# Epigenetic clock and lifespan prediction in the short-lived killifish *Nothobranchius furzeri*

**DOI:** 10.1101/2024.08.07.606986

**Authors:** Chiara Giannuzzi, Mario Baumgart, Francesco Neri, Alessandro Cellerino

## Abstract

Aging, characterized by a gradual decline in organismal fitness, is the primary risk factor for numerous diseases including cancer, cardiovascular, and neurodegenerative disorders. The inter-individual variability in aging and disease susceptibility has led to the concept of “biological age,” an indirect measure of an individual’s relative fitness. Epigenetic changes, particularly DNA methylation, have emerged as reliable biomarkers for estimating biological age, leading to the development of predictive models known as epigenetic clocks. Initially created for humans, these clocks have been extended to various mammalian species. Here we set to expand these tools for the short-lived killifish, *Nothobranchius furzeri*. This species, with its remarkably short lifespan and expression of canonical aging hallmarks, offers a unique model for experimental aging studies.

We developed an epigenetic clock for *N. furzeri* using reduced-representation bisulfite sequencing (RRBS) to analyze DNA methylation in brain and caudal fin tissues across different ages. Our study involved generating comprehensive DNA methylation datasets and employing machine learning to create predictive models based on individual CpG sites and co-methylation modules. These models demonstrated high accuracy in estimating chronological age, with a median absolute error of 3 weeks (7.5% of median lifespan) for a clock based on methylation of individual CpG and 1.5 weeks (3.7% of median lifespan) for an eigenvector-based clock.

Our investigation extended to assessing epigenetic age acceleration in different strains and the potential resetting effect of regeneration on fin tissue. Notably, our models indicated that a shorter-lived strain has accelerated epigenetic aging and that regeneration does not reset, but may decelerate epigenetic aging. Additionally, we used longitudinal data to develop an “epigenetic timer” for direct prediction of individual lifespan based on fin biopsies and eigenvector-based method, achieving a median absolute error of 4.5 weeks in the prediction of actual age of death. This surprising result underscores the existence of intrinsic determinants of lifespan established early in life.

This study presents the first epigenetic clocks and lifespan predictors for *N. furzeri*, highlighting their potential as aging biomarkers and sets the stage for future research on life-extending interventions in this model organism.

## Introduction

Aging is the gradual and progressive decline of fitness with advancing age and constitutes the main common risk factor for cancer, cardiovascular-, and neurodegenerative-diseases (Stambler et al., 2018). Within aging populations, considerable inter-individual variability exists in disease susceptibility and mortality risk, prompting the conceptualization of “biological age” to denote an individual’s relative fitness within an aging cohort (Jackson et al., 2003). Given the challenge of directly measuring relative fitness, indirect methods to estimate “biological age” have been proposed. Age-dependent molecular changes in cells and tissues represent quantifiable proxies of organismal fitness. Notably, epigenetic alterations, a canonical hallmarks of aging and age-related diseases (Issa, 2014; Rakyan et al., 2010; Teschendorff et al., 2010), stand out as reliable proxy of age. Specifically, certain CpG sites in the genome exhibit consistent age-dependent methylation patterns and represent highly informative input variables for developing predictive models of biological age, termed ‘epigenetic clocks’.

Initially developed as quantitative biomarkers of human aging (Horvath, 2013; Levine et al., 2018; Lu et al., 2019; Belsky et al., 2020), epigenetic clocks have been gradually extended to other mammalian species. The conservation of age-dependent methylation of some specific CpGs across the mammalian class (Horvath, 2013; Horvath et al., 2020; Wang et al., 2017a, 2020) enabled the construction of an universal pan-tissue epigenetic clocks capable of predicting age of 9 tissue types from 128 different mammalian species (Lu et al., 2023). The association of CpGs informative for age prediction with developmental genes (Lu et al., 2023) suggests an evolutionary conserved mechanism linking aging and mammalian development, although these consistent alterations do not appear to be causally implicated in aging (Ying et al., 2024).

Teleost fishes are increasingly acknowledged as valuable models for investigating disease mechanisms (Beck et al., 2022; Schartl, 2013) and aging (Benjamin, 1965; Carneiro et al., 2016; Gerhard & Cheng, 2002; Ghoneum & Egami, 1982; Liu & Walford, 1966; Markofsky & Milstoc, 1979; Poeschla & Valenzano, 2020) motivating the development of epigenetic clocks for these taxa. An investigation of age-dependent DNA methylation changes in the teleost *Nothobranchius furzeri* would be of particular interest. This species has adapted to survive in the ephemeral ponds of the African savannah where their survival is constrained by the seasonal monsoon. In captivity, their lifespan is limited between 3 and 8 months (Terzibasi et al., 2008; Valdesalici & Cellerino, 2003), the shortest recorded among vertebrates. This compressed lifespan is associated to rapid acceleration of age-dependent mortality, explosive growth, rapid age-dependent functional decline and expression of canonical hallmarks of aging (Cellerino et al., 2016; Hu & Brunet, 2018) including increased activity of the polycomb repressive complex (Baumgart et al., 2014; Cencioni et al., 2019) an hallmark of the regions associated with mammalian epigenetic clocks (A. T. Lu et al., 2023; Moqri et al., 2022). While epigenetic clocks have been established for different teleost species (Anastasiadi & Piferrer, 2020; Mayne, Espinoza, et al., 2021; Mayne et al., 2020; Piferrer & Anastasiadi, 2023), none currently exists for *N. furzeri*. Here, we set to develop an accurate and generalizable epigenetic clock for this species using a sequencing-based approach.

Epigenetic clocks in humans and most mammals rely on DNA methylation data generated on Illumina Infinium microarrays (Hannum et al., 2013; Horvath, 2013; A. T. Lu et al., 2023). By contrast, recent methylation studies in other organisms relied on high-throughput bisulphite sequencing (BS-seq) approaches (e.g. Chatterjee et al., 2013; Chen et al., 2015; Hahn et al., 2017; Pegoraro et al., 2016; Platt et al., 2015; Stubbs et al., 2017; Wang et al., 2017b), for two main reasons. The development costs of species-specific microarrays are prohibitive, leading to a preference for sequencing-based methodologies. Secondly, sequencing-based approaches offer the flexibility to explore differential methylation patterns beyond the regions targeted by DNA methylation arrays (W. Zhang et al., 2015; Y. Zhang et al., 2016). Here, we established a protocol for DNA methylation analysis through reduced-representation bisulphite sequencing (RRBS), a genome-wide approach at single-nucleotide resolution that combines restriction enzyme digestion, DNA size selection and bisulfite sequencing to enrich for areas of the genome with a high CpG content, where DNA methylation is concentrated (Smith & Meissner, 2013). However, RRBS suffers from an intrinsic limitation, namely the limited overlap in detected CpGs across independent experiments, probably due to the enzymatic reactions involved in the library construction that result in variable genome coverage across experiments (Field et al., 2018b; Thompson et al., 2018). This variability hinders the transfer of epigenetic clocks to datasets outside the original study. To develop a more generalizable epigenetic model we used as input co-methylation modules (Lu et al., 2023) that enable correlation-based imputation of missing values. Leveraging a cross-sectional dataset, we developed models that can accurately predict the age of *N. furzeri* from brain and caudal fin tissues. Next, we applied a longitudinal design were we obtained fin biopsies of fish at young age, recorded their age at death and applied these methods to investigate the relationship of epigenetic age with mortality. Deviations of epigenetic age from calendar age, known as epigenetic age acceleration, serve as markers of diseases (Fransquet et al., 2019) but, as accuracy in age prediction increases, association with mortality risk vanishes. Notably, epigenetic clocks predicting risk factors (GrimAge, PhenoAge) (Levine et al., 2018b; A. T. Lu et al., 2019, 2022; Q. Zhang et al., 2019) or the rate of physiological decline (DunedinPACE) (Belsky et al., 2022) have shown superior capabilities of predicting mortality risk. To train a mortality predictor in *N. furzeri*, we exploited the longitudinal information and trained a model to predict age at death directly. Finally, we investigate whether the fin regeneration process caused a resetting of the epigenetic age in the regenerated tissue utilizing a subset of the same cohort

## Materials and methods

### Animals and rearing conditions

#### Fish maintenance

*Nothobranchius furzeri* fish (strains MZM0410 and GRZ) were bred and maintained in the fish facility of the Leibniz Institute on Aging – Fritz Lipmann Institute (Jena, Germany) as described in Baumgart et al. (2014). Fish were housed individually in 3L plastic tanks with divider in a recirculating system (Aqua Schwartz, Göttingen) on a 12-h light/12-h dark cycle, water temperature 26°, water conductivity 2.5 mS and were fed once a day with alive *Chironomus* larvae. Fish condition was monitored daily by trained animal caretakers and fish were euthanized if one of the endpoints of the severity assessment was reached. Euthanasia was performed by rapid chilling.

#### Longitudinal sample collection

For all experimental fish, a fin biopsy was removed from the upper tip of the caudal fin with a scalpel under anesthesia (MS-222) at age 10 weeks. The biopsy was further divided into two parts, one was used in the present study and the second was stored for future studies. For all fish that survived until 20 weeks, a second fin biopsy was removed from the lower tip of the caudal fin taking care to exclude regenerated tissue of the upper caudal fin. All biopsies were performed at 10 am in fasted state, to avoid effects of circadian rhythms and feeding status. Samples were immediately placed on dry ice and stored in an ultralow freezer (−80°C) until DNA extraction.

#### Ethical statements

Fish were reared in the facility according to §11 of German Animal Welfare Act under license number J-003798. The animal experiment protocols were approved by the local German authority in the State of Thuringia (Veterinaer-und Lebensmittelueberwachungsamt; longitudinal study: reference number 22-2684-04-FLI-17-001).

#### Datasets

We generated 5 datasets, analyzing, in total, n=418 *Nothobranchius furzeri* tissue samples from 2 different tissues, 2 strains, both male and female, spread over 13 different ages. “Dataset 1” was built in a cross-sectional experiment using 12 male fish of the strain MZM-0410: brain and caudal fin were collected from each fish at the stages of sexual maturity (5 weeks, n=4), young adult (13 weeks, n=4) and old (38 weeks, n=4). “Dataset 2” included caudal fin samples collected from 15 male and 31 female fish, all of the strain MZM-0410, of ten different ages. “Dataset 3” was obtained using 24 tail fin biopsies from 15 male and 9 female fish of the strain GRZ, collected at six different ages. “Dataset 4” were obtained from a longitudinal experiment, in which caudal fin was taken from 153 fish, all male and of the strain MZM-0410, at 10 weeks and at 20 weeks of age. The lifespan was recorded for all 153 fish. “Dataset 5” was derived using 5 animals from the longitudinal study with which Dataset 4 was generated: before these fish were sacrificed for reaching a human endpoint, the piece of fin that had grown back after trimming at 10 or 20 weeks of age was recut.

### DNA extraction

DNA was isolated from snap-frozen tissue using the QIAamp® DNA Micro Kit (Qiagen) following the manufacturer’s protocol. DNA concentration was measured with NanoDrop 2000 (Thermo Scientific), using 1 μl of input DNA. Then, dsDNA concentration was quantified using Quant-iT Picogreen dsDNA Assay (Thermo Scientific) and the distribution of DNA fragments was assessed by TapeStation 4200 (Agilent). The quality criteria to submit the samples to sequencing were: DNA Integrity (DIN) > 8, A260/280 ratio between 1.8 and 2.0 and A260/230 ratio above 2.0.

### Reduced representation bisulfite sequencing (RRBS) method

RRBS libraries were produced in the DNA Sequencing Facility of the FLI by Ovation RRBS Methyl-Seq with the TrueMethyl oxBS kit (Tecan, Redwood City, CA, USA) according to the manufacturer’s instructions. Briefly, 100 ng of genomic DNA was digested for 1h at 37 °C with the methylation-insensitive restriction enzyme MpsI. Then, fragments were ligated to methylated adapters and treated with bisulfite to convert unmethylated cytosine into uracil. PCR amplification was then performed to obtain the final DNA library. The resulting libraries were sequenced on the Illumina NovaSeq6000 with 100 bp single-end sequencing with an average of reads per sample of: 180.6 ± 42.3 million for Dataset 1, 58 ± 5,9 mln for Datasets 2 and 3 that were processed together, 82.7±6.3 mln for Dataset 4 and 59,3 ± 5,8 mln for Dataset 5.

### RRBS data analysis

#### Alignment and data preprocessing

The sequencing reads were trimmed to remove adapter and low-quality bases with Trim Galore! (http://www.bioinformatics.babraham.ac.uk/projects/trim_galore/). Then, using the program BSMAPz (https://github.com/zyndagj/BSMAPz) trimmed reads were aligned to the *Nothobranchius furzeri* MZM-0410 reference genome. This was generated by combining the *N. furzeri* GRZ reference genome fasta file with the MZM variants vcf file (both downloaded from the *Nothobranchius furzeri* Genome Browser, https://nfingb.leibniz-fli.de/), via the FastaAlternateReferenceMaker tool from GATK. Using the python script methratio.py in BSMAPz, methylation ratios were extracted from the mapping output. Methylation analysis was performed using the R package *methylKit* (version 1.20.0) (<u>Akalin et al., 2012</u>). A filter based on read coverage was applied: bases with coverage below 10X were discarded and bases in the 99.9th percentile of coverage in each sample were discarded, in order to eliminate PCR artifacts. The methylation ratio of each site was calculated by dividing number of reads calling “C” by the total number of reads calling either “C” or “T” at the specific site. The methylation score for each CpG site is represented as a β-value which ranges from 0 (unmethylated) to 1 (fully methylated). Differentially methylated CpGs (DMCpGs) were detected using a logistic regression model (in *methylkit*) based on a Chi-square test. A false discovery rate (FDR) q value threshold of < 0.05 and methylation difference of ±5% between case and control groups was used to identify significant DMCpGs.

#### Principal component analysis (PCA)

A principal component analysis (PCA) was used to visualize sample clustering based on the global variation in CpG methilation. PCA is a dimensionality-reduction method that projects the original samples into a lower-dimensional space based on the correlation pattern among input variables and it is useful to reveal data structure and influence of variables such as tissue and age on global methylation patterns. PCA was performed using the *prcomp()* function that uses the singular value decomposition in order to examine the correlations between individuals, and then the R package *factoextra* to extract and visualize its output.

#### Annotation of mCpG genomic context

To test for CpGs enrichment among transcriptional features, the file ‘Nfu_20150522.genes_20150922.gff3’ (downloaded from the *Nothobranchius furzeri* Genome Browser, https://nfingb.leibniz-fli.de/) was used to define genomic regions as promoters, introns, exons, or intergenic. Promoter was defined as the ±1kb region surrounding the transcription start site. The CpG sites were then annotated to one of these regions using the *annotateWithGeneParts()* function from the R package *Genomation*. When CpG sites mapped to more than one genomic region, the following precedence was applied: promoter>exon>intron. To annotate CpG loci also at CpG islands, shores and flanking regions, a CpG islands track was constructed on the MZM-0410 reference sequence using the *cpg_lh* program from the UCSC Genome Browser, and used in the *annotateWithFeatureFlank()* function, similar to the previous one. Gene annotation was performed only for those CpGs that fell into promoters or gene bodies, the *N. furzeri* genes were then mapped to the human orthologues by BLAST. This procedure detected 12258 human orthologues of 18980 *N. furzeri* genes that were then annotated with the corresponding human gene name.

#### Enrichment analysis of CpG associated genes

Enrichment analysis was conducted using the web tools WebGestalt (https://www.webgestalt.org/). Two databases were selected: gene ontology (GO) database, which contains information about molecular function, biological process, and cellular component, and KEGG pathway database that has a collection of known pathways in each of the genes involved. When multiple CpGs were annotated to the same gene, the gene was counted only once. For all enrichment tests, the background was all the genes containing CpG sites passing the filter criteria described in “Alignment and preprocessing”. To select and rank the significantly enriched GO terms and pathways a false discovery rate (FDR) < 0.05 was used.

### Development of a generalizable epigenetic clock

We employed the R package *caret* (version 6.0-94) (Kuhn, 2008) to establish two epigenetic clocks for estimating chronological age across two tissues (brain and fin) in MZM-0410 *N. furzeri* fish (n=70, Datasets 1 and 2). The two models differed in type of input used:

1. Individual CpG sites (clock 1)
2. Eigenvectors of CpG co-methylation modules (clock 2)

#### Site-based clock

We used methylation values of single CpG sites as input, selecting those covered in all 70 samples with a coverage ≥ 10X and in the 99.9^th^ percentile. Samples were randomly assigned to a training- (80%, n=56) or test-set (20%, n=14) using the createDataPartition function in the *caret* R package. To limit the inclusion of invariant or uninformative sites, we filtered CpGs in the training set according to their correlation with age using the cpg.assoc function in the eponymous R package (FDR <0.05). The model was trained taking advantage of the R package *caretEnsemble* that allows a comparison of the performance of several user-defined regression algorithms. Specifically, we selected eleven methods: Independent Component Regression (icr), Support Vector Machines with Radial Basis Function Kernel (svmRadialSigma), Partial Least Squares Generalized Linear Model (plsRglm), Partial Least Squares Models (widekernelpls, kernelpls, simpls and pls), a rule-based model (cubist), generalized linear model via penalized maximum likelihood (glmnet), random forest (rf), and elastic net (enet). Each of these methods performs feature selection independently as part of the model training to reduce the number of input variables, so to reduce the risk of overfitting and improve the generality of the models. All the eleven regression models were trained with the same resampling parameters: three repeats of 10-fold cross-validation, used to further mitigate overfitting. We selected the elastic net model (enet) based on its overall performance (R^2^, RMSE and MAE) and evaluated its predictive capability on the test set (20% of Datasets 1 and 2), by regressing all predicted age values onto known chronological ages.

#### Eigenvector-based clock

Machine learning algorithms are prone to overfitting and show better performances with reduced numbers of predictor variables (Badillo et al., 2020). We reduced the number of input features by identifying clusters (or modules) of CpGs with high correlation in methylation and summarizing methylation of each cluster by its methylation eigenvector. Using the R package *WGCNA* (Langfelder & Horvath, 2008), we computed a weighted correlation network on the set of CpGs covered in all the 70 samples by at least 10 reads and filtered based on age correlation with the function cpg.assoc. We applied an FDR threshold of 0.2 on the correlation of methylation with age in the train set to remove redundant sites and sites with missing values, but retaining a suitable number of sites to enable network generation. We chose a suitable soft-thresholding power (sft, power β=12) using the pickSoftThreshold function that evaluates the conformity of the co-expression network with a scale-free topology. Then, we employed the function blockwiseModule to automatic detect modules via dynamic tree cutting, using the parameter deepSplit 2. For each module, we summarized the CpG methylation levels with its first eigenvector (eigengene). Thus, we used all eigenvectors as input variables and adopted the same pipeline with the same parameters as the site-based clock to train and validate the second algorithm. To further improve the model performance, here we not only compared the individual regression methods, but also combined them with the function caretStack in the *caretEnsemble* R package, resulting in a meta-model with superior performances than each of the individual models.

### Model Validation with Leave-One-Out cross validation (LOOCV)

We evaluated the stability of the models with two cross validation strategies: leave-one-fraction-out (LOFO) and leave-one-sample-out (LOSO). For LOFO, the 70 samples from Dataset 1 and Dataset 2 were split into ten balanced and evenly sized fractions. LOSO and LOFO cross validation trained each model on all but one sample or fraction, respectively, using three repeats of 10-fold internal cross validation. The left-out sample/set was then used as a test set and its predicted age was saved. When iterations were completed, a predicted age value was available for each sample of the dataset. In this analysis, we used the same CpGs/eigenvectors/tiles selected for building the respective clock. The R packages *caret* and *caretEnsemble* were used with the same input features, parameters and regression models as described above for the three different models. Once all age predictions were generated (independently for each of the three clocks), the predicted age values were regressed onto known chronological ages to evaluate the performance of the method.

### Model Validation in Independent Datasets

We investigated selected biological aspects involved in the aging process by applying model 2 to three independently generated RRBS datasets (Dataset 3, Dataset 4 and Dataset 5). The direct applicability of model 1 was hindered by the limited overlap between the independent datasets. For clock 2 it was necessary to re-train the model on all CpGs present in both Datasets 1 and 2 and test set (Datasets 3 or 5, separately). For GRZ biopsies (Dataset 3) the sites covered in at least 80% of the samples were retained for downstream processes. As samples of Dataset 3 were processed and sequenced together with Dataset 2, batch effects can be excluded, which is not the case for Dataset 5 whose samples were processed separately. Therefore, for the latter, a new model was created to mitigate the effects of the batch on the prediction. Dataset 4 is close to Dataset 5 in PCA space, so a group of 70 random samples from Dataset 4 was included in the training set together with samples from Datasets 1 and 2. The sites covered in at least 90% of the samples from the 140 in the training set and those to be tested from the regenerated fins and respective longitudinal study samples derived from the same fish were selected. The same model was also used to test the remaining samples from Dataset 4 that were not included in the training set. Then, for all three test sets, the same data processing workflow developed above was used, building a weighted CpG correlation network through which input variables could be obtained to train a new eigenvector-based model.

### Development of an epigenetic timer

The eigenvector-based model was used to directly estimate remaining lifespan, generating what we call an “epigenetic timer”. To this end, we leveraged of Dataset 5 for which the age of death of the individual fish is reported in addition to the methylation data. First, CpG sites detected in all 306 samples of Dataset 5 were selected and then, using the varFilter function in the *genefilter* R package (v.1.76.0, 2021), a statistical variance filter was applied to remove features exhibiting little variation across samples (var.cutoff=0.77). Using a soft-thresholding power of 14, network construction and module detection were performed (deepSplit=3). Modules that were closely related based on similarity measured by the correlation of their eigenvectors were merged (cutHeight=0.6). Samples were randomly divided into training (80%, n=246) and test set (20%, n=60) using the createDataPartition function in the *caret* R package. The R package *caretEnsem* trained the final model using *ble* to compare 13 regression algorithms at the same time: Independent Component Regression (icr), Support Vector Machines with Polynomial Kernel (svmPoly), Support Vector Machines with Radial Basis Function Kernel (svmRadialSigma), Partial Least Squares Generalized Linear Model (plsRglm), Gaussian Process with Polynomial Kernel (gaussprPoly), Partial Least Squares Models (widekernelpls, kernelpls, simpls and pls), a rule-based *model* (cubist), generalized linear model via penalized maximum likelihood (glmnet), random forest (rf), and elastic net (enet). Training was performed with three repeats of 10-fold cross-validation. Analysis of overall performance (R^2^, RMSE and MAE) selected the ‘gaussprPoly’ method and it was then applied on the 20% of samples left out of the training set to evaluate its performance in predicting fish lifespan. The predicted values were thus regressed onto ages at death to evaluate model fit by calculating R^2^, the root mean square error (RMSE) and the mean absolute error (MAE).

### Model Validation with Leave-One-Out cross validation (LOOCV)

The accuracy of our epigenetic timer was evaluated with two cross validation methods, as for models predicting age: leave-one-out (LOO) and leave-one-set-out (LOSO). For LOSO, the 306 samples from Dataset 5 were assigned to ten evenly sized sets. To performed both LOO and LOSO cross validation, one sample/set was held out at a time from the regression, and a model was trained on the remaining N-1 sample/set, using three repeats of 10-fold internal cross validation. The left-out sample/set was then used as a test set. Following the pipeline described above, using the R packages *caret* and *caretEnsemble*, the model was trained with the above-mentioned 13 regression algorithms, either in combination with each other or specifically with the gaussprPoly method only, selected in the previous step. Once all age predictions were generated, regression of the predicted lifespan values onto the actual lifespan provided an assessment of the model performance.

## Results

### Age-Dependent DNA Methylation in *N. furzeri* brain and fin

To characterize the patterns of age-dependent DNA methylation in *N. furzeri*, we initially sequenced at high coverage a total of 24 libraries obtained from samples of caudal fin and brain of 12 male animals of the strain MZM0410 at the life stages of sexual maturity (5 weeks), young adult (13 weeks) and old (38 weeks ~ median lifespan). We name this dataset “Dataset 1” (Table 1).

**Tabella 1.**
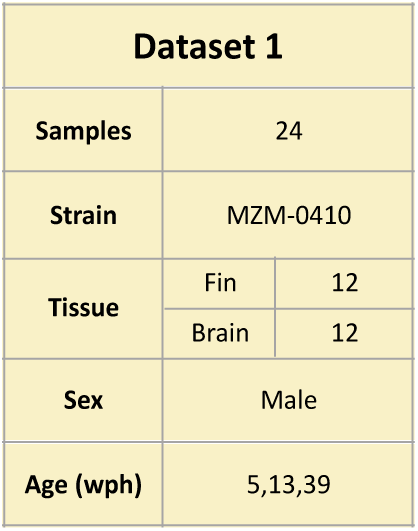

We obtained an average of 180.6 million reads per sample (range 58.1-284.4) that correspond to 2,398,438 CpG covered by at least 10 reads in all samples that were retained for downstream analysis (Fig.1A-B). After filtering, we obtained 2,392,796 CpG covered in all samples. The distribution of the percentage of methylation revealed the expected bimodal distribution with most of the CpGs present at the extremes of the distribution (Fig.1C). The distribution of genomic location of the reads showed the enrichment in promoter regions and CpG islands expected for RRBS data (Fig.1D-E) (Guo et al., 2015; Sun et al., 2015).

**Figure 1.**
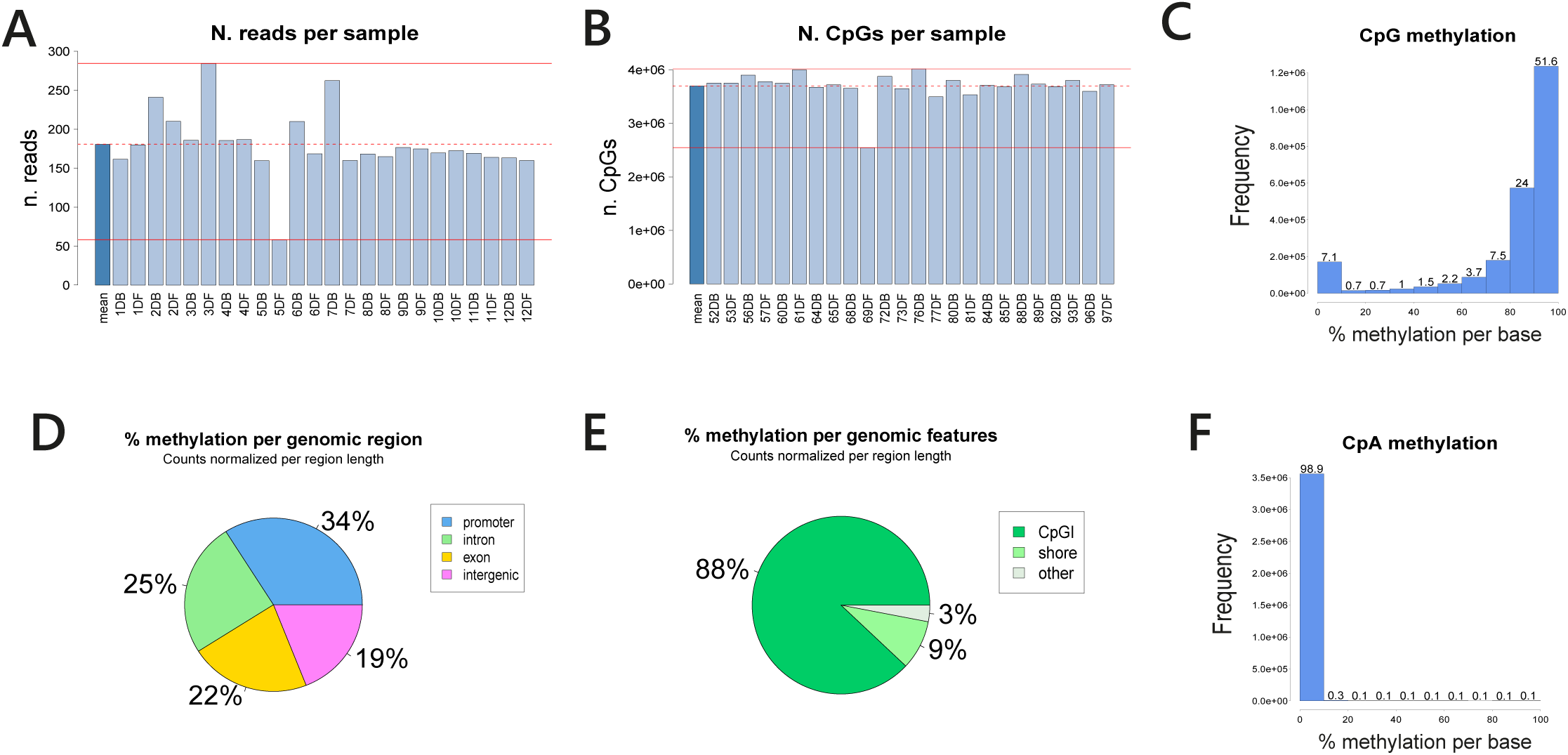
Quality control and genomic features of sampled CpGs in Dataset 1. (**A**) Number of reads per sample. Red lines indicate the minimum and maximum values, the dotted line represents the mean value. (**B**) Number of CpGs covererd per sample. Red lines indicate the minimum and maximum values, the dotted line represents the mean value. (**C**) Distribution of CpG methylation fraction of the filtered CpGs. (**D**) Distribution of genomic regions for the sampled CpGs. (**E**) Distribution of genomic for the sampled CpGs. CpGI stands for CpG island and shore for CpG shore. (**F**) Distibution of CpA methylation.

We estimated the bisulphite conversion rate by calculating the methylation level of cytosines outside the CpG context. We calculated the level of methylation in CH context (where H corresponds to A, T or C). Methylation of CA is a prominent feature of DNA methylation in mammalian brains (Lister et al., 2013; Ramsahoye et al., 2000; Ziller et al., 2011). Therefore, cytosines in CA and CT context were analyzed and we could not detect methylation, confirming the excellent conversion efficiency of bisulphite in our experiment (Fig.1F) and highlighting a difference between mammalian- and fish-brains in the context of CA methylation.

In order to assess global similarities between young, adult and old fish methylomes across the two tissues, all CpGs obtained after pre-processing were used as input variables for principal component analysis (PCA). PCA clearly separated the samples according to their groups (Fig.2A). The first component (PC1, 53.5% of variance explained) separated samples according to tissues and the second component (PC2, 5.5% of variance explained) discriminated the three age groups of either tissue.

**Figure 2.**
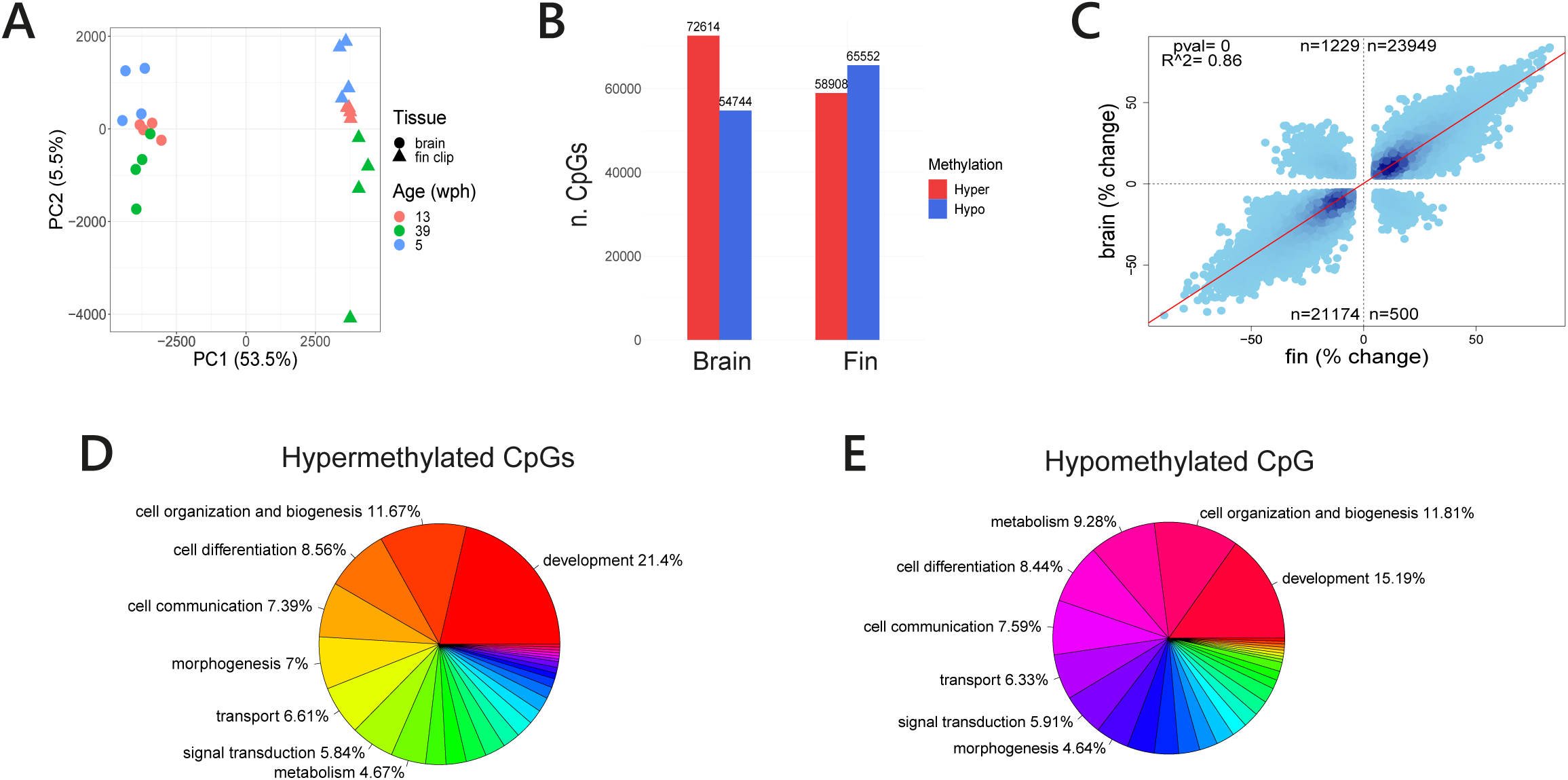
Age dependent regulation of CpG methylation in Dataset 1. (**A**) PCA of all the samples. Color codes for age and shape for tissue type: triangle for fin and circle for brain. (**B**) Number of hyper- and hypo-methylated CpGs with age in brain and fin. (**C**) Scatterplot of methylation changes with age in brain and fin. Only CpGs significant in both tissues are reported. (**D**) Distribution of GO terms in the genes associated with positively age-regulated CpGs. (**E**) Distribution of GO terms in the genes associated with negatively age-regulated CpGs.

We then investigated the effect of age on DNA methylation by calculating differentially methylated CpGs (DMCpGs) between old and young fish for the two tissues separately using the R package ‘methylKit’, applying a Chi-square test. This analysis revealed that, after selecting CpGs with a methylation difference greater than 5% and a q-value < 0.05, 72,614 CpGs were hypermethylated and 54,744 CpGs were hypomethylated with age in the brain and 58,908 CpGs were hypermethylated with age and 65,552 CpGs were hypomethylated with age in the fin (Fig.2B). Direct comparison of the two tissues revealed that 46,852 CpG are differentially methylated with age in both brain and fin, of which 96.3% are regulated in a coherent direction in both tissues (Fig.2C).

Further, we analyzed the DMCpGs located within or in close vicinity of an annotated gene, focusing on those coherently up- (quadrant 1, Fig.2C) or down-regulated (quadrant 3, Fig.2C) in both tissues. 14,724 hypermethylated CpGs (61.5%) and 12,765 hypomethylated CpGs (60.3%) could be linked to an annotated gene. A Gene Ontology (GO) overrepresentation analysis of those genes revealed a significant enrichment of gene associated with development and specifically development of the nervous system both in hyper-and hypo-methylated CpGs (Fig.2D-E and Supplementary Table 1).

### Development and validation of epigenetic clocks in *N. furzeri*

To generate an epigenetic clock for *N. furzeri*, we sequenced 46 additional libraries of fin samples from male and female MZM0410 individuals. We selected fin as target tissue because it enables longitudinal sampling in living fish and our previous results show that its methylation dynamics mirrors brain methylation dynamics. We selected samples spanning an age range from 3- to 44-weeks in order to cover the lifespan of this strain evenly and produced “Dataset 2” (Table 2).

**Tabella 2.**
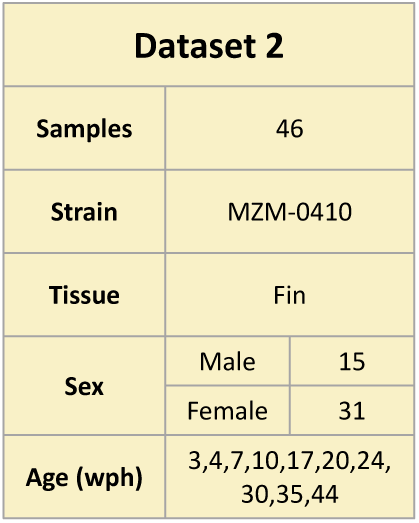

We aimed at obtaining a shallower coverage of a larger number of samples and obtained an average of 58 ± 5.9 million reads per sample that correspond to 307,652 CpG covered by at least 10 reads in all samples that we retained for downstream analysis. After filtering, we obtained 304,596 CpG covered in all samples.

The total number of samples in Dataset 1 and Dataset 2 was 70, which corresponds to the estimated minimum number of samples necessary to generate an epigenetic clock in fish (Mayne, Berry, et al., 2021). Brain samples from Dataset 1 were included as their age-dependent methylation closely resembles the methylation in the skin. The number of CpGs detected in Dataset 1 and present also in Dataset 2 was 294,175, corresponding to 15.1% of the CpGs detected in Dataset 1 and 96.6% of those detected in Dataset 2.

We used two approaches in order to identify the most informative input variable for model building. In the first (clock 1, site-based), we used as input DNA methylation level of individual CpGs, covered in all 70 samples. To reduce input dimensionality and facilitate model training, we split the data in train- (80%) and test-set (20%) and reduced the number of predictor variables to 90, filtering those with significant correlation with age in the train set (FDR < 0.05). We compared the performances of different regression models using the R package *caret* and found elastic net to be the most accurate (R^2^ = 0.95) with a median absolute error (MAE) of 3 weeks between chronological age ad DNA methylation-based age estimate in the test set (Fig.3A - clock1). We recalculated the data splitting into train- and test-set 10 times, using the same partitioning percentages, but a different seed for each iteration. This approach allowed the model to randomly select a different small subset of CpG sites each time, each capable of accurately predicting the chronological age of the animals (Supplementary Table 2). Additionally, we retrained the model using an increasing number of CpG sites, each time selecting the sites from a list ranked on correlation with age. The results indicated that 90 CpGs provide the highest predictive accuracy, with an improvement from 30 to 90 sites and a progressive decline from 90 to 900 (Supplementary Table 3).

**Figure 3.**
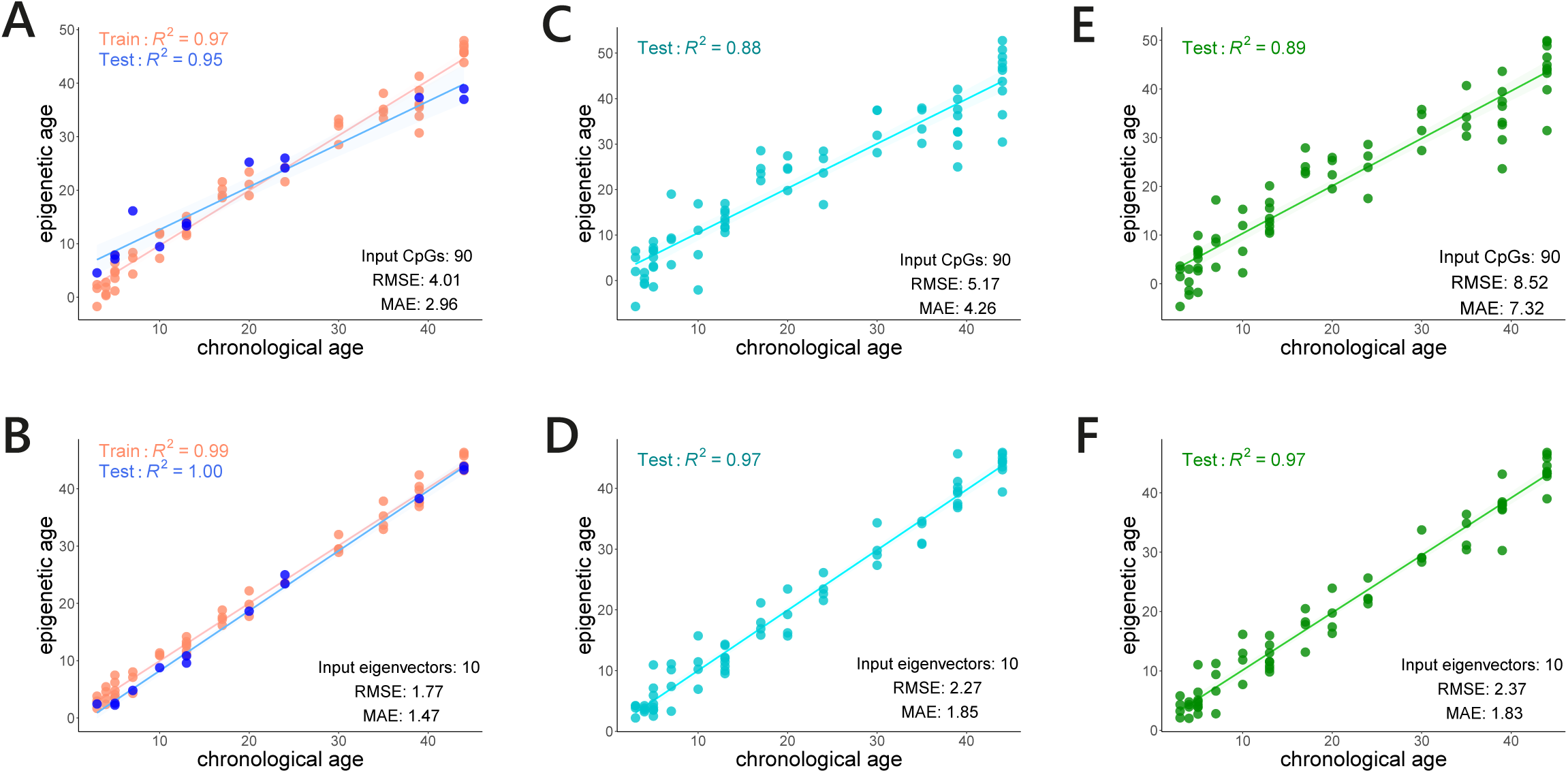
Epigenetic clock for *N. furzeri* displayed as scatter plots. Real age is reported on the abscissa and predicted age on the ordinate. (**A**) Age prediction in the train- and test set of site-based clock 1. (**B**) Age prediction in the train- and test set of eigenvector-based clock 2. (**C**) Leave one fraction out validation of clock1. (**D**) Leave one fraction out validation of clock2. (**E**) Leave one sample out validation of clock1. (**F**) Leave one sample out validation of clock2. Correlation values and error metrics are reported as insets in all plots.

We developed a second clock (clock 2) to reduce the number of input variables further leveraging on the high level of correlation among individual CpGs by building a weighted gene co-expression network (WGCN) of methylated CpGs. We identified 10 modules of co-regulated CpGs and for each of these methylation modules we calculated the eigenvector as a summary variable for the coherent methylation pattern within the module and used these as input for model training. We refer to clock 2 as eigenvector-based clock. Here, a combination of the 11 models selected in the R package *caret* produced a better prediction than the single best regression model and this meta-model provided an age predictor with an average error of only 1.5 weeks in the test set and an age correlation approaching 1 (Fig.3B - clock2). We therefore deployed an ensemble of algorithms instead of a single one also in the following cross-validation steps.

To evaluate the stability of our epigenetic clocks, we employed two canonical testing methods: leave-one-fraction-out cross-validation (LOFO-CV) and leave-one-sample-out cross-validation (LOSO-CV). LOFO-CV divided the 70 samples (Dataset 1 and Dataset 2) into ten sets and each served as test set in one iteration, while the remaining nine fractions constituted the training set. LOSO-CV excluded one sample instead of one fraction per iteration leaving all the others as test set. The strong correlation between chronological- and predicted-age persisted in the cross-validation models: R^2^ = 0.88 and 0.89 for clock 1, R^2^ = 0.97 and 0.97 for clock 2 for LOFO- and LOSO-CV, respectively (Fig.3C-F). This evaluation also established clock 2 as the most accurate predictor of age.

### Relationship between H3K27 methylation and DNA methylation

Aging-associated epigenetic remodeling are not limited to DNA methylation, but extend to histone posttranslational modification such as H3K27me3 (Booth & Brunet, 2016a; Brunet & Berger, 2014). The latter contributes to heterochromatin formation and gene silencing, particularly of development-related genes (Cai et al., 2021). We took advantage of a published dataset of young and old *Nothobranchius furzeri* skeletal muscle samples in which the density of H3K27me3 modification was quantified using chromatin immunoprecipitation (ChIP) sequencing (Cencioni et al., 2019) and investigated the relationship between H3K27 methylation and DNA methylation in our RRBS dataFare clic o toccare qui per immettere il testo.. We found that among all 294,175 CpGs detected in our Datasets 1 and 2, only 11% were found in regions of significant H3K27me3 peaks, in contrast with 46% of the 90 CpG sites selected by the site-based clock, demonstrating that predictive CpGs are enriched for H3K27me3 peaks (Fig.4A, Fisher’s exact test, p-value = 1.25e-16).

**Figure 4.**
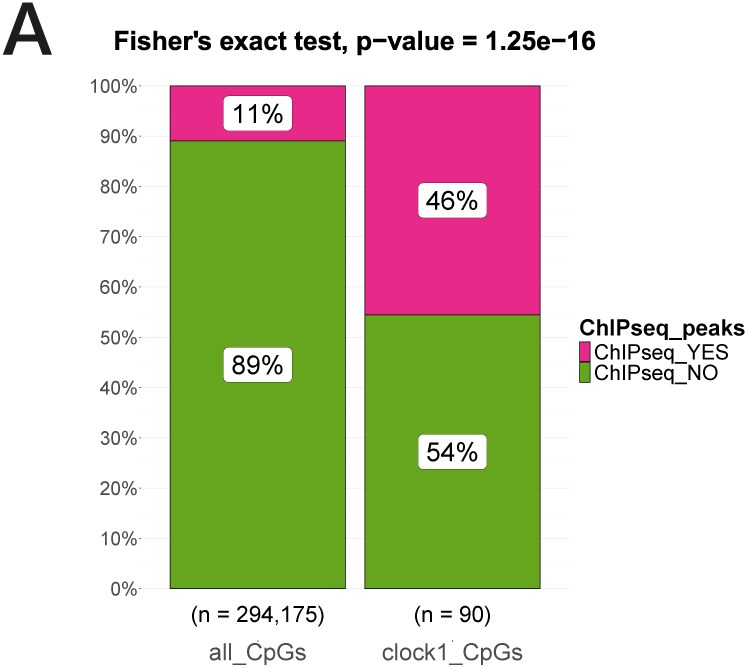
Correspondence between H3K37me3 peaks and CpGs used to construct clock1. Green indicates overlap and purple lack of overlap.

### Assessing epigenetic age acceleration in a short-lived *N. furzeri* strain

Different strains of *N. furzeri* show large heritable differences in lifespan (Terzibasi et al., 2008). To build our epigenetic clocks, we used the strain MZM0410 with an average lifespan of 40 weeks. On the other hand, the GRZ strain with a median lifespan of 20 weeks in our colony (Fig.5A). This short lifespan is associated with accelerated expression of cellular age-related phenotypes and accelerated increase of age-dependent mortality (Terzibasi et al., 2008). We applied our clocks to Dataset 3, obtained from 24 fins of GRZ fish of both sexes (Table 3) to test epigenetic age acceleration in this strain. Only 1,267 CpGs were common between the 70 samples of the training set (Datasets 1 and 2) and all the 24 of the test set (Dataset 3), highlighting the problem of generalizability of site-based clocks. To avoid filtering of all CpGs with one or few missing values and maximize information retained, we selected the sites covered in at least 80% of the samples (n = 26,896 CpGs), used a correlation-based method of imputation of missing values and retrained an eigenvector-based clock. We observed for clock 2 R^2^ = 0.99 for the MZM0410 train set an overall decent correlation between the chronological age and the best fit of predicted age: R^2^ = 0.75 from GRZ (Fig.5B). Reduced accuracy could result from prominent strain-specific methylation patterns as revealed by principal component analysis (Fig.5C), which showed that GRZ samples form a separate cluster. This could explain the difficulty in generalizing a model trained with MZM0410 to GRZ. The model captures the underlying differences in biological ageing across strains, as the short-lived GRZ strain consistently shows a significantly higher DNA methylation-based age than the longer-lived MZM0410 strain, demonstrating that epigenetic age acceleration correlates with shorter lifespan and higher aging rate of GRZ.

**Figure 5.**
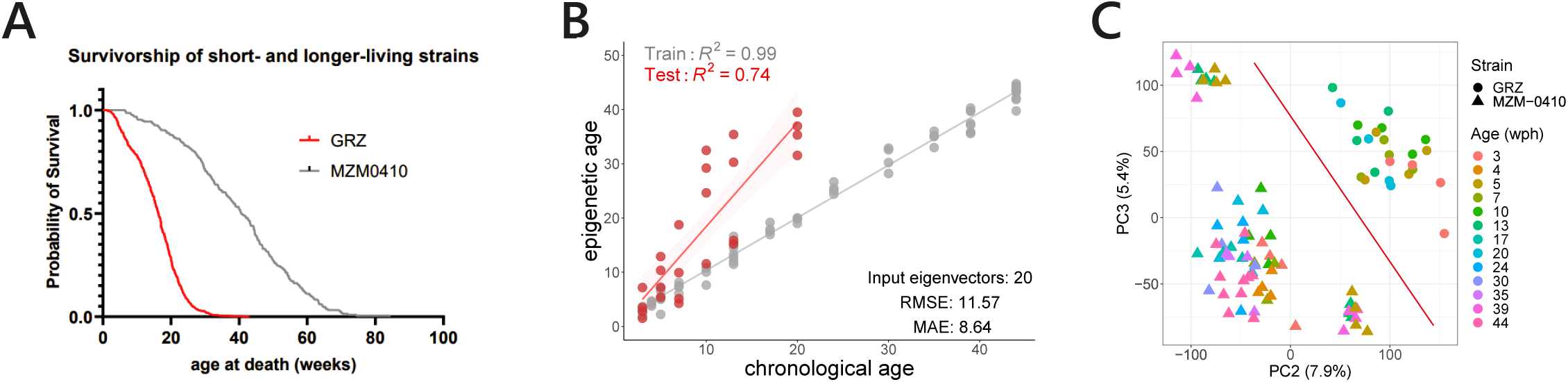
Epigenetic clock validation in the short-lived strain GRZ. (**A**) Lifespan of the strain GRZ and MZM0410 in the FLI colony. (**B**) Age prediction as scatter plots. The model was trained on MZM0410 and tested on GRZ. Real age is reported on the abscissa and predicted age on the ordinate. Correlation values and error metrics are reported as insets the plot. (**C**) PCA of the samples belonging to the two different strains.

**Tabella 3.**
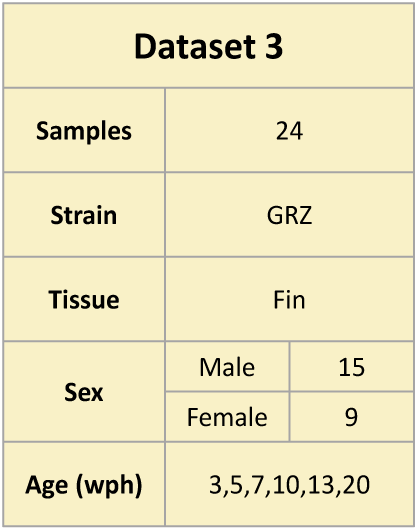

### Epigenetic age and lifespan in *N. furzeri*

To test whether epigenetic age acceleration is a predictor of short lifespan and whether regeneration influences epigenetic age, we collected caudal fin biopsies at 10 and 20 weeks of age and recorded time to death longitudinally from 153 fish (Dataset 4: Table 4 and Fig.7A). Principal component analysis (PCA) revealed a pronounced batch effect with a separation of Dataset 4 along the first component (PC1, 58.6% of variance explained) from those of Datasets 1 and 2, which were used to create the models (Fig.6A). This is an expected complication when analyzing RRBS experiments sequenced at different times. Since the intersection of CpGs covered in Datasets 1, 2, 4 and 5 was small (n = 1,613), given the large number of samples included, we selected those covered in at least 90% of the samples (n = 18,683) to calculate the input eigenvectors for clock 2 and retrained the model by adding 70 randomly selected samples from Dataset 4 to the existing 70 samples from Datasets 1 and 2 in the training set and tested the remaining samples of Dataset 4. We then used this retrained model to calculate the epigenetic age of samples from Dataset 4 not included in the training set. Using the WGCN generated in Fig.6B, we calculated the eigenvector of each module with the methylation values of respective CpGs in Dataset 4 samples. This new model provided a good prediction of age with a median absolute error (MAE) of almost 7 weeks (Fig.6C). To examine whether the individual deviation of epigenetic age from real age (epigenetic acceleration) is associated with lifespan, we regressed epigenetic age against lifespan at both ages using as predictor variable epigenetic age at 10 and 20 weeks separately (Fig. 6D), the average of these two values (Fig. 6E) or the slope between 10- and 20-weeks (Fig. 6F). None of these analyses detected a significant correlation between epigenetic age acceleration and age at death.

**Figure 6.**
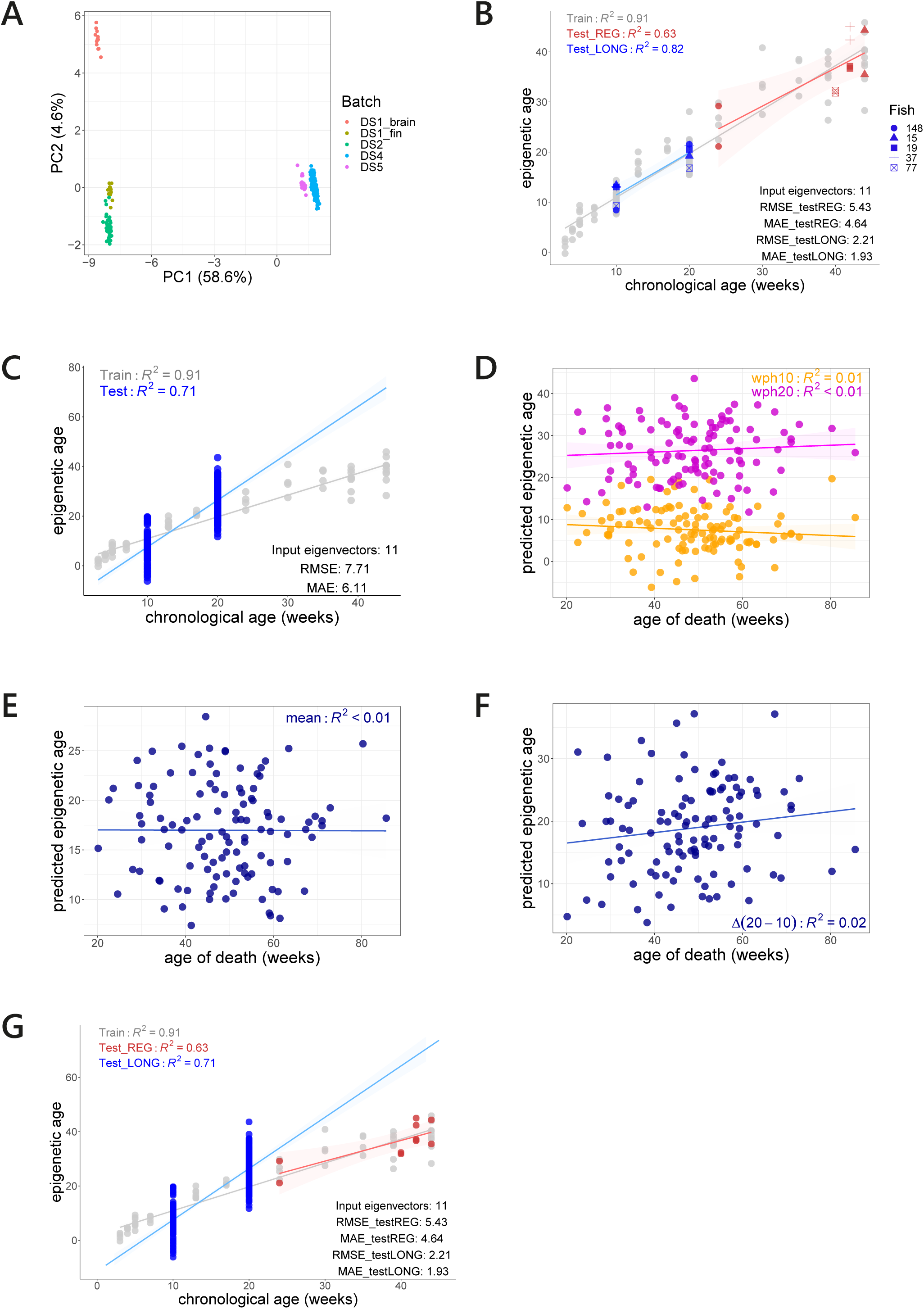
Epigenetic age of regenerated fin. (**A**) PCA of samples belonging to the different datasets. The color indicates the different dataset and, for Dataset 1, the different tissues. (**B**) Performance of the age predictor retrained including 70 random samples from the longitudinal study displayed as scatter plots. For each sample, real age is reported on the abscissa and predicted age on the ordinate. The shape of the points corresponds to different donors. Blue points are the two fin samples taken before regeneration and the red points correspond to two samples of the regenerated part of the same fin. (**C**) Age prediction of the samples from the longitudinal study not used for retraining the epigenetic clock. (**D-F**) Relationship between epigenetic age acceleration and actual lifespan. (**D**) Correlation separated on the age at collection of the samples: orange points correspond to samples collected at 10 weeks and purple points to samples collected at 20 weeks. (**E**) Average of correlations of the same donor obtained from samples collected at 10- and 20-weeks. (**F**) Delta of correlations of the same donor obtained from samples collected at 10- and 20-weeks. (**G**) Overlapping age predictions of samples from the longitudinal study tested in Fig.6B (blue points) and regenerated fin samples tested in Fig.6C (red points).

**Figure 7.**
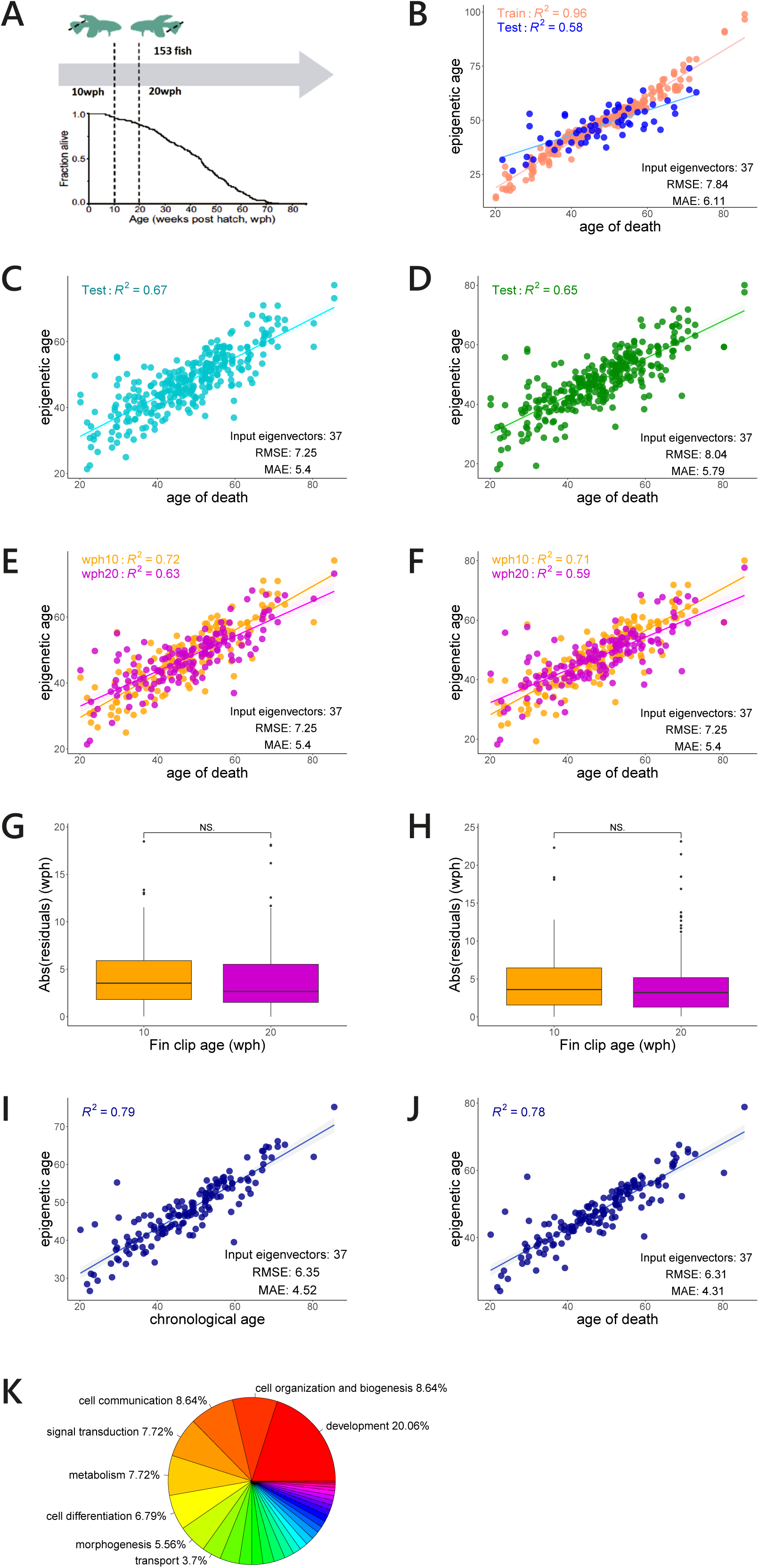
Epigenetic lifespan predictor. (**A**) Scheme of the longitudinal study. (**B-F**) Performance of the lifespan predictor displayed as scatter plots. For each sample, real age at death of the donor is reported on the abscissa and predicted age at death on the ordinate. (**B**) Lifespan prediction in the train- and test set of eigenvector-based timer. (**C**) Leave one fraction out cross-validation validation. (**D**) Leave one sample out validation. (**E-F**) Predictions separated on the age at collection of the samples: 10 weeks represented as orange dots and 20 weeks as purple dots. (**E**) Leave one fraction out cross-validation validation. (**F**) Leave one sample out cross-validation. (**G-H**) Error distribution for predictions separated on the age at collection of the samples: 10 weeks represented as orange boxplot and 20 weeks as purple boxplot. (**G**) Leave one fraction out cross-validation validation. (**H**) Leave one sample out cross-validation. (**I-J**) Predictions per individual obtained by averaging the predictions of the same donor obtained from samples collected at 10- and 20-weeks. (**I**) Leave one fraction out cross-validation validation. (**J**) Leave one sample out cross-validation. (**K**) Distribution of GO categories in the eigenvectors used for model construction.

**Tabella 4.**
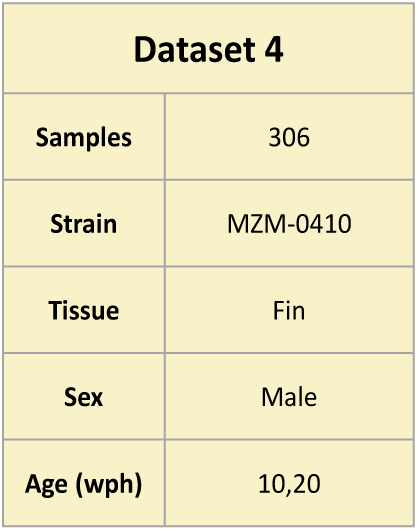

### Epigenetic age of regenerated tissue

To test the effect of regeneration on epigenetic age, we analyzed Dataset 5 that contains regenerated fin clips from a subset of fish from the longitudinal study euthanized for humane reasons (Table 5). Samples of dataset 5 cluster close to samples of Dataset 4 in PCA space (Fig. 6A). Using the same model built to calculate the epigenetic age of Dataset 4 samples, we were able to test epigenetic age of the new set of regenerated fins, mitigating the influence of batch on the prediction (Fig.6B). As a further test of the effective removal of batch effects, we also included as test sets the respective samples from Dataset 4 derived from the same fish before regeneration at 10 and 20 weeks. The epigenetic age of the regenerated tissue increased with the age of the fish, indicating that regeneration does not reset epigenetic age. However, overlaying the predictions for Dataset 4 samples shown in Fig.6C with those for Dataset 5 samples in Fig.6B, shows that the regenerated fin samples points all fall below the regression line that connects the mean predicted epigenetic ages for the two groups within Dataset 4, suggesting a deceleration of age-dependent DNA methylation in the regenerated tissue (Fig.6G).

**Tabella 5.**
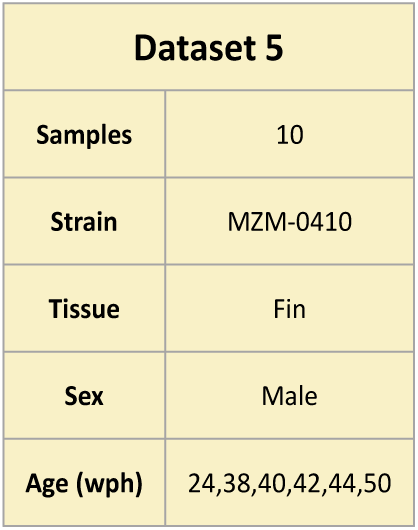

### Direct prediction of age at death

Previous human studies indicate that models trained to predict age are poor predictors of mortality risk as opposed to models directly trained on risk factors or aging rate (Levine et al., 2018; A. T. Lu et al., 2019; Belsky et al., 2020). We therefore trained a predictor directly on age at death and developed an “epigenetic timer”. Since the eigenvector-based clock 2 provides the most accurate age prediction, we developed a model starting with 89,763 methylation sites common among the 306 fin samples in Dataset 4, we filtered the 20.646 sites of highest variance to build a weighted CpG correlation network (Langfelder & Horvath, 2008) and identified 37 modules of co-methylated CpGs whose eigenvector served as input variable for training the predictor. We split the dataset into a training and a test set (80% and 20% respectively) and demonstrated that this eigenvector-based timer can predict fish lifespan. The combination of the 13 selected regression methods allowed for achieving a correlation between estimated and actual age of death (R^2^) of 0.58 and MAE of 6.11 weeks for the best partition (Fig.7B). To obtain unbiased estimates of the model performance, we validated it by the two cross-validation methods described above, (LOFO-CV and LOSO-CV), obtaining MAE of 5.4 and 5.8 weeks, respectively (Fig.7C-D).

The model provides two separate predictions for the same fish based on the input data collected at 10- and 20-weeks separately. We thus compared the accuracy of these two predictions. Surprisingly, we found no significant difference in the accuracy of these predictions assessed by correlation (Fig.7E-F) or absolute error (Fig.7G-H). On the other hand, accuracy improves when the two predictions are averaged (LOFO-CV and LOSO-CV, R^2^ = 0.79 and 0.78, MAE = 4.5 and 4.31 weeks, respectively Fig.7I-J). Overall, these results suggest that individual variability in lifespan is strongly influenced by intrinsic conditions that are established in early adulthood.

To characterize biological functions associated with the CpGs informative for prediction of time to death, we performed a Gene Ontology (GO) overrepresentation analysis (ORA) of genes in which the CpGs belonging to modules selected by our epigenetic timer fell. Similarly to what previously observed for age prediction, a significant enrichment of genes associated with development was detected (Fig.7K and Supplementary Table 4).

## Discussion

Here, we present the first epigenetic clock for *N. furzeri*. We used RRBS and quantified methylation level of several hundred thousand CpGs from two tissue types (brain and fin) and 13 different ages for a total of 70 samples of both sexes distributed over a range from 3 weeks (before sexual maturity) to 44 weeks (past median lifespan) in the longer-lived strain MZM0410 and 7 different ages for a total of 25 samples for the shorter-lived GRZ strain. Further, we generated a longitudinal dataset comprising two fin biopsies of 153 fish in order to correlate DNA methylation with individual lifespan. These two datasets represent a unique resource for *N. furzeri* and for comparative analysis of age-dependent DNA methylation across Vertebrates. Using these DNA methylation data, we identified pathways associated with epigenetic aging in *N. furzeri* and developed the first epigenetic age- and lifespan-predictors for this model.

### Genomic features associated with age-dependent DNA methylation in *N. furzeri*

A detailed analysis of CpGs differentially methylated with age revealed several salient features of the epigenetic aging in this species. First, the vast majority (96.3%) of sites differentially methylated with age in both fin and brain are coherently hyper- or hypo-methylated, suggesting that also in the killifish aging is associated with a coordinated DNA methylation signature across many organs as described in mammals (A. T. Lu et al., 2023).

Analysis of GO terms overrepresentation among genes in proximity of CpGs hypermethylated with age showed maximum enrichment for GO terms related to developmental processes and particularly nervous system development. This finding is consistent with a similar enrichment of hypermethylated CpGs in developmental genes detected in humans and in a wide range of mammals (Horvath, 2013; A. T. Lu et al., 2023; Stubbs et al., 2017; Thompson et al., 2017) Fare clic o toccare qui per immettere il testo.highlighting a remarkable conservation of age-related CpG methylation dynamics across Vertebrates. Development is characterized by epigenetic remodeling via DNA methylation that irreversibly restricts cell fate and prevents dedifferentiation (Reik et al., 2001). The preferential methylation of genes most active during development lends support to the view of quasi-programmed aging. According to this view, aging results from the failure in adult life to completely halt regulatory programs specific of earlier phases of ontogeny (Blagosklonny, 2006; Booth & Brunet, 2016b; Bowles, 1998; Gems, 2022; Horvath & Raj, 2018; Magalhães, 2012; Mayne et al., 2019; Mitteldorf, 2016; Rando & Chang, 2012; Sen et al., 2016; J.-H. Yang et al., 2019).

Ample evidence indicates that age-associated DNA methylation is more likely to occur in regions targeted by the Polycomb repressive complex in humans and more generally in mammals (Horvath, 2013; Horvath et al., 2012; Jasinska et al., 2022; A. T. Lu et al., 2023). Polycomb repressive complexes are involved in regulation of gene expression via histone H3 lysine 27 trimethylation (H3K7me3), inducing compaction and gene silencing. Specifically, PRC2 represses key developmental gene promoters needed to establish a specific cell lineage (Cai et al., 2021). Age-dependent methylation is particularly enriched in bivalent promoters where both repressive H3K27me3 and permissive H3K4me3 modifications are deposited (Rakyan et al., 2010). A relationship between H3K27me3 and DNA methylation is evident during development. In embryonic stem cells, H3K27me3 usually occurs in unmethylated regions (Brinkman et al., 2012; Statham et al., 2012). During differentiation, DNA methylation occurs in promoters that were marked with H3K27me3 in embryonic stem cells (Mohn et al., 2008) as the result of a possible sequential process by which silencing is initially established by polycomb repressive complexes in early development and is later consolidated by long term by DNA methylation. Therefore, persistent increase of DNA methylation with age represents a continuation of this developmental process. Alternatively, it was recently shown that hepatocytes lose H3K27me3 peaks with age, but these are replaced by DNA methylation, explaining the age-dependent methylation of these regions (N. Yang et al., 2023).

### Two different models to compute killifish epigenetic age

Mammalian epigenetic clocks for mammals are usually array-based. This technology cannot be easily adapted to teleost fishes. In order to develop our predictors in *N.furzeri*, we made use of RRBS, which is a general and a cost-effective technique to quantify DNA methylation at single base resolution that takes advantage of restriction enzymes and generates a reduced representation of the genome enriched in CpG islands and promoter regions. On the backside, the reduced representation represents also a major limitation of the RRBS method, as the genome coverage varies from experiment to experiment resulting in batch effects and limited overlap between the CpGs sequenced in independent experiments. As a consequence of this variability, the model (clock) trained on given datasets may not be applied directly to an external dataset because some of the predictive CpGs will not be present in this second dataset.

One further challenge in the development of ML predictors based on epigenetic data is the disparity between number of input CpG sites and number of samples in the training set. The high dimensionality of the input dataset lends to redundancy and overfitting. For this reason, we employed different strategies to reduce the dimensionality of the predictor variables. In the site-based clock 1, in which the predictor variables were methylation levels of individual CpGs, we followed Horvath (2013) and filtered for CpGs whose methylation level in the training set is significantly correlated with age. A second approach followed Lu et al., (2023) and took advantage of high levels of correlation between different CpGs to reduce the number of input variables, grouping CpGs with a similar profile into co-methylation modules whose first eigenvector was used as predictive variable (eigenvector-based clock).

During the construction of the different epigenetic clocks, we noticed that, although each CpG site has a different degree of association with age, the set of CpGs that can be used collectively to accurately estimate age is large. Indeed, using different seeds at each iteration of training, site-based clocks randomly selected a different small set of CpGs, each supporting an accurate prediction of age. This finding is consistent with the observation of redundancy in predictive CpGs in the human epigenome that enables the construction of many different aging clocks from the same DNA methylation data (Porter et al., 2021). Epigenome aging is connected to developmentally driven processes but also entails a stochastic component leading to the accumulation of random errors in the epigenome, often described as “epigenetic drift” (López-Otín et al., 2013; Martin, 2005). This global stochastic process may partially explain the redundancy of the aging clocks and very recent evidence indicates that stochasticity in DNA methylation can be used as input variable to predict age (Meyer & Schumacher, 2024; Tong et al., 2024).

### Modulation of epigenetic age

We applied the predictors we developed to examine how epigenetic age is modulated in some relevant biological contexts. We first investigated age-dependent DNA methylation in the short-lived GRZ strain. A first result is that genotype has a larger influence on DNA methylation that age itself, with samples from the two strains showing clear separation in PCA space despite being sequenced in the same batch. Age prediction in GRZ was less accurate as compared to MZM0410, likely due to the idiosyncratic epigenetic patterns in the GRZ strain. The GRZ strain showed significant epigenetic age acceleration as compared to the longer-lived strain MZM0410. These two results, large strain-dependent differences and accelerated transcriptomic aging in the shortest-lived strain, are congruent with previous analysis of RNA-seq data (Mazzetto et al., 2023) and indicate that faster aging rate is associated with epigenetic age acceleration in GRZ.

The analysis of epigenetic age in regenerated fins was faced with confounding batch effects. Therefore, we retrained the model including some samples of Dataset 4 (longitudinal study) in the training set. These new model correctly predicted the age of fin samples and was used to test Dataset 5 (regenerated fins). These initial results on effects of regeneration on epigenetic age indicate that regeneration does not reset epigenetic age, but may slow the accumulation of age-dependent epigenetic changes. The data derived from samples of the longitudinal study obtained in a limited number of individuals and our conclusions should be considered as preliminary. These initial data, however, warrant future specifically-designed studies to investigate epigenetic age modulation in regenerated tissue of killifish.

Finally, we tested whether epigenetic age was predictive of lifespan in a prospective study. We could not detect an association between epigenetic age and mortality, in line with the result of human studies where epigenetic age obtained from clock trained to predict chronological is a poor predictor of mortality when compared to model trained to predict risk factors (GrimAge, PhenoAge) or the rate of physiological decline (DunedinPACE).

### An epigenetic timer predicts lifespan

The same machine learning approach used to construct the age predictor was set to estimate the time-to-death in *N. furzeri* leveraging on a longitudinal experimental design. Our epigenetic “timer” uses as input the methylation levels at 10- and 20-weeks of age. Since epigenetic drift accumulates over time, a better prediction of time to death for models trained on information collected at an older age is expected. In contrast, we observed no significant difference in prediction accuracy from data collected at these two time points, suggesting that conditions limiting lifespan are established early in life at least in *N. furzeri* under laboratory conditions. When information from the two ages is combined, individual lifespan prediction reached R^2^ of around 0.8 i.e., 80% of lifespan variation is accounted for by the epigenetic timer. This striking result probably stems from the highly controlled conditions under which these fish are raised that minimize variations in environmental and dietary factors and unravel intrinsic determinants of lifespan, highlighting the value of *N. furzeri* for longitudinal studies.

### Applications as biomarkers

Interest in epigenetic clocks originates mostly from their possible use as biomarkers for intervention studies in animal models and humans. They are envisaged as a method to test dietary and pharmacological interventions for their putative life-extending action without the need to perform life-long experiments that have lifespan as the primary endpoint of analysis. *N. furzeri* represents a particularly convenient vertebrate model for lifespan intervention studies (Astre et al., 2023; Baumgart et al., 2016; Moses et al., 2024; Ripa et al., 2023; Valenzano et al., 2006). The epigenetic clocks and timers we developed could therefore represent valuable tools to accelerate intervention studies in *N. furzeri.* For the moment, these models were trained in the long-living strain MZM0410 and we observed a large epigenetic difference between this strain and shorter-lived GRZ strain reducing cross-strain transferability of the models. Future developments will therefore prioritize the training of epigenetic clocks in the GRZ as well as a longitudinal study in this shorter-lived strain. A final validation of these tools will require to test the effects of established life-extending manipulations on both clock and timer to test whether they are sensitive to interventions. If these validation experiments will provide a positive result, then epigenetic biomarkers of aging for the killifish will become very valuable tools to enhance the use of this species as model organism for investigations of life-extending interventions.

## Acknowledgments

We thank Cinzia Caterino per excellent technical assistance. We thank the Fish Facility of the Leibniz Institute on Ageing, Fritz Lipmann Institute for fish breeding and husbandry and the DNA Sequencing Facility of the Leibniz Institute on Ageing, Fritz Lipmann Institute for the generation of the primary sequencing data.

